# Genetic contribution to two factors of neuroticism is associated with affluence, better health, and longer life

**DOI:** 10.1101/146787

**Authors:** W David Hill, Alexander Weiss, Andrew M McIntosh, Catharine R Gale, Ian J Deary

## Abstract

Neuroticism is a personality trait that describes the tendency to experience negative emotions. Individual differences in neuroticism are moderately stable across much of the life course^1^; the trait is heritable^2-5^, and higher levels are associated with psychiatric disorders^6-8^, and have been estimated to have an economic burden to society greater than that of substance abuse, mood, or anxiety disorders^9^. Understanding the genetic architecture of neuroticism therefore has the potential to offer insight into the causes of psychiatric disorders, general wellbeing^10^, and longevity. The broad trait of neuroticism is composed of narrower traits, or factors. It was recently discovered that, whereas higher scores on the broad trait of neuroticism are associated with earlier death, higher scores on a ‘worry/vulnerability’ factor are associated with living longer^11^. Here, we examine the genetic architectures of two neuroticism factors—worry/vulnerability and anxiety/tension—and how they contrast with the architecture of the general factor of neuroticism. We show that, whereas the polygenic load for general factor of neuroticism is associated with an increased risk of coronary artery disease (CAD), major depressive disorder, and poorer self-rated health, the genetic variants associated with high levels of the anxiety/tension and worry/vulnerability factors are associated with affluence, higher cognitive ability, better self-rated health, and longer life. We also identify the first genes associated with factors of neuroticism that are linked with these positive outcomes that show no relationship with the general factor of neuroticism.

---

Participants in the present study were members of the UK Biobank sample (http://www.ukbiobank.ac.uk)^12^. All analyses included only those individuals who self-described as White British. We analysed Genome-Wide Association Study (GWAS) single nucleotide polymorphism (SNP) data that passed quality control from 91,469 participants (mean age 56.75 years, 47,246 females) who completed the 12 neuroticism questions from the Eysenck Personality Questionnaire-Revised Short Form^13^. We used these data to derive a general factor of neuroticism from the 12 items. Next, two factors were extracted from the residual variance one of these factors we called anxiety/tension, and the other worry/vulnerability, both of which were independent of, or orthogonal to, the general factor (Online Methods, **Supplementary Table 1**). The general factor correlated with the anxiety/tension factor at *r* = 0.069, and with the worry/vulnerability factor at r = 0.121, (*P* = < 2.20 × 10^−16^), and the factors correlated with each other at r = 0.427 (*P* = < 2.20 ×10^−16^). The general factor of neuroticism correlated phenotypically at *r* = 0.957 (*P* = < 2.20 × 10^−16^), and genetically at *r*_*g*_ = 0.978, (SE = 0.005) with scores on the full neuroticism scale, indicating that it is almost equivalent to the neuroticism measure published by Smith et al.^2^ In the current study we include the general factor of neuroticism to show how the specific neuroticism factors of anxiety/tension and worry/vulnerability not only have a different genetic architecture, but in contrast to the general factor, are associated with a polygenic load for better cognitive and physical health. All three phenotypes were adjusted for age, gender, assessment centre, genotyping batch, genotyping array, and 14 principal components in order to correct for population stratification prior to all analyses (**Supplementary Figure 1**.).

We estimated the heritability of each of the neuroticism phenotypes using GREML conducted in GCTA. A total of 14.6% (SE = 0.7%) of the phenotypic variation of the general factor of neuroticism is explained by the additive effects of common genotyped SNPs. This is comparable to the original estimate by Smith et al.^2^. SNP-based heritability for each of the specific neuroticism factors are reported here for the first time with the additive effects of common genotyped SNPs explaining 7.8% (SE = 0.7%) of the variance in the anxiety/tension factor, and 9.7% (SE = 0.7%) of the worry/vulnerability factor. The general factor of neuroticism was found to have a genetic correlation with the anxiety/tension factor, *r*_*g*_ = 0.312 (SE = 0.073, *P* = 1.85 × 10^−5^), and with the worry/vulnerability factor, *r*_*g*_ = 0.293, (SE = 0.070, *P* = 2.76 × 10^−5^); the two factors had a strong genetic correlation with each other, *r*_*g*_ = 0.646, (SE = 0.052, *P* = 4.87 × 10^−36^).

Genome-wide association analyses for the general neuroticism factor and the two factors were performed using an imputed dataset that combined the UK10K haplotype and 1000 Genomes Phase 3 reference panels; details of the phasing and imputation procedure can be found at http://biobank.ctsu.ox.ac.uk/crystal/refer.cgi?id=157020. Univariate LD regression conducted on the each GWAS indicated that there was no residual stratification left after correcting on the 14 principal components (**Supplementary Table 2**.), and that the inflation in test statistics was due to the presence of a polygenic signal rather than confounding.

For the general factor of neuroticism we identified 1,436 SNPs that were genome wide significant and formed 11 independent loci. Seven of these were on chromosome 8, two on chromosome 18, one on chromosome 9, and one on chromosome 2 (**Figure 1. Table 1 & Table 2**). Again, these findings were comparable to those from the original study by Smith et al.^2^. We include them here in order to compare them with the first GWASs of neuroticism factors, which we report next.

**Figure 1.**
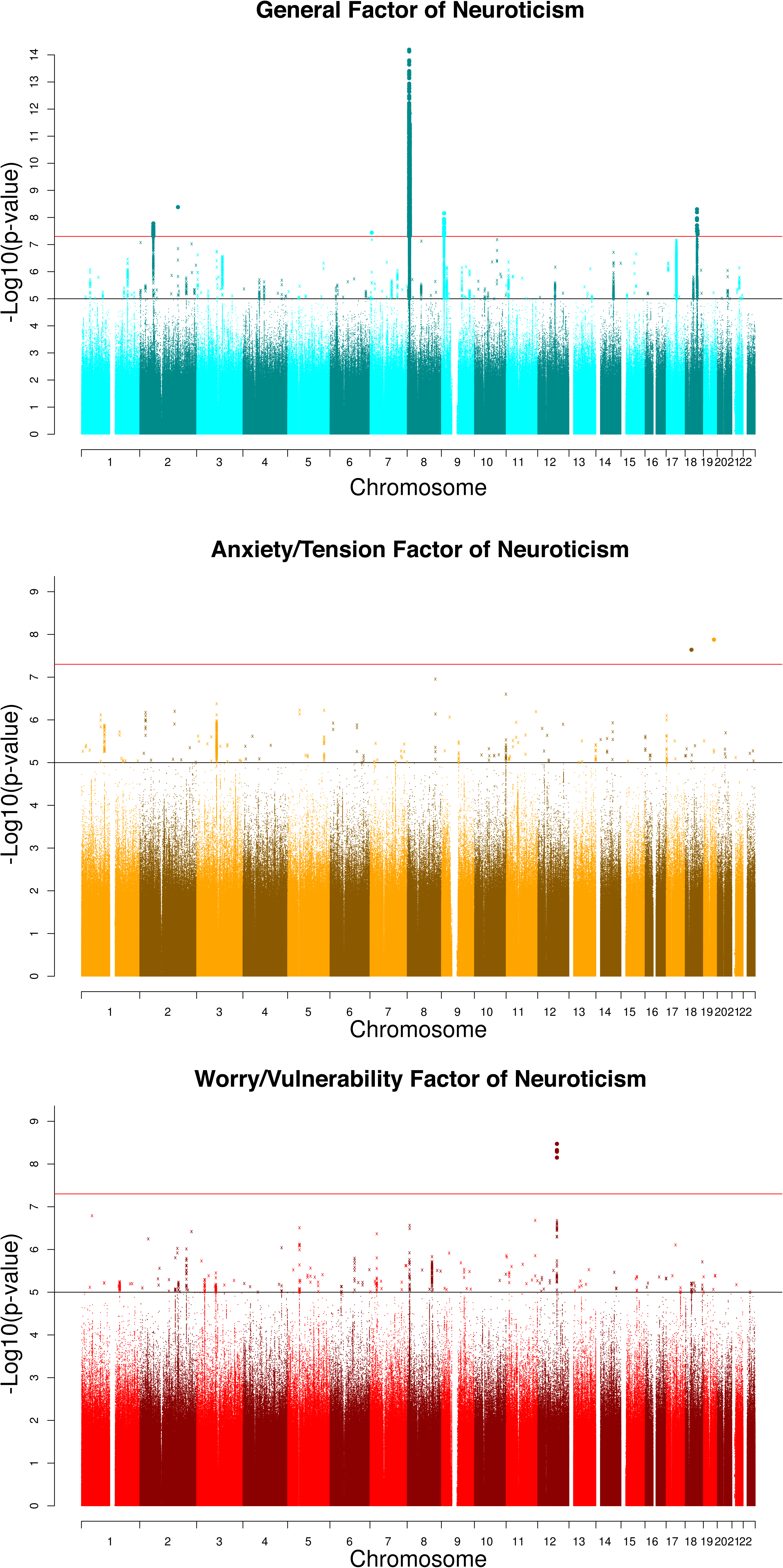
Manhattan plots for the general factor of neuroticism (Blue), the anxiety/tension factor (Orange), and the worry/vulnerability factor (Red). All samples sizes were 91,469 participants. The red line indicates genome wide significance.

**Table 1.**
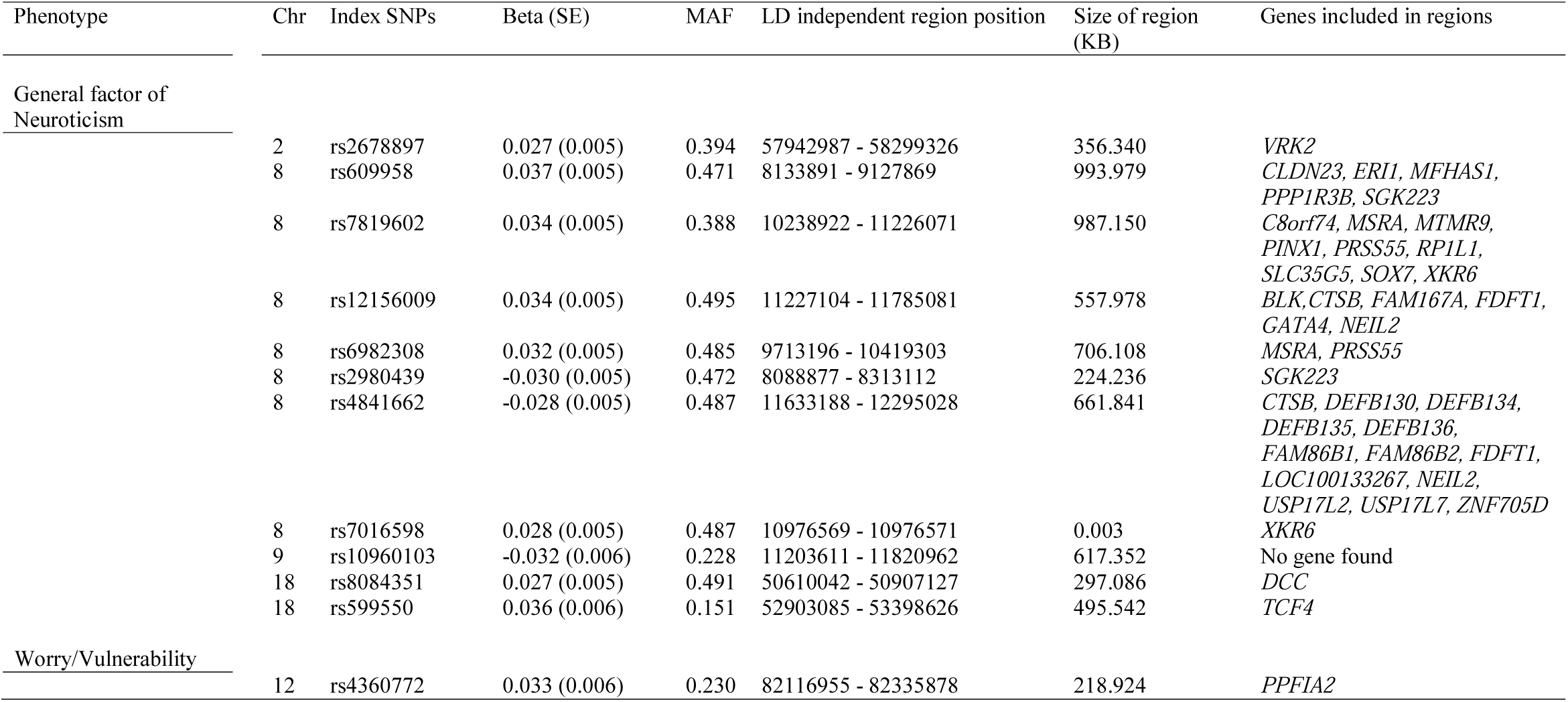
Genome wide significant index SNPs for the General factor of Neuroticism and the two factors of Anxiety/Tension, and Worry/Vulnerability. Index SNP indicates the most significant SNP n each LD clump. No regions were identified using the LD clumping method for the anxiety/tension factor.

**Table 2.**
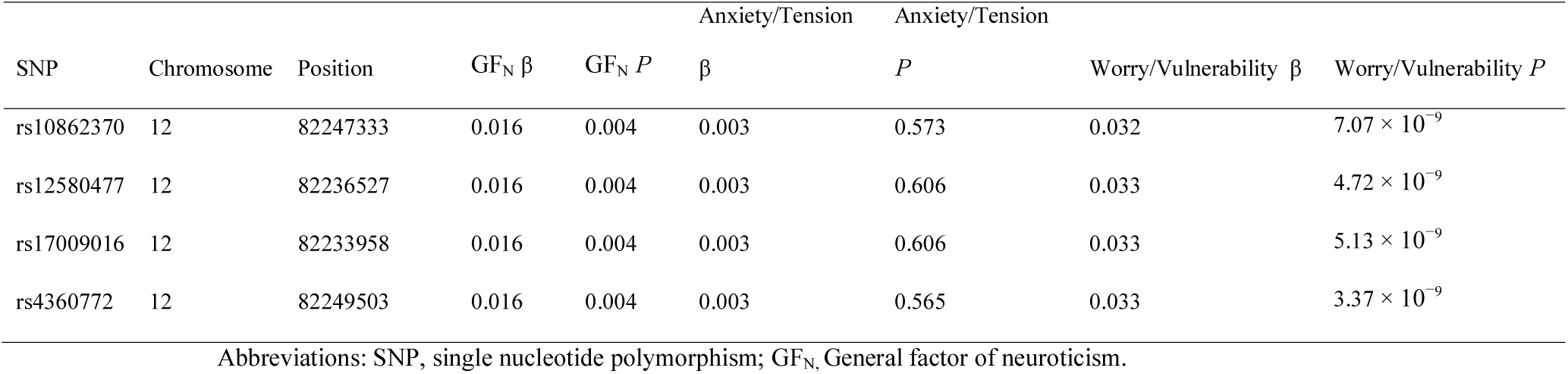
Comparing the genome-wide significant SNPs from the worry/vulnerability phenotype with the General factor of neuroticism and the anxiety/tension factor.

**Table 3.** Comparing the genome-wide significant genes in the anxiety/tension, and the worry/vulnerability phenotype with the general factor of neuroticism. Significant genes are highlighted in bold.

| Gene | CHR | START | STOP | Worry/Vulnerability $P$ | Anxiety/Tension $P$ | General factor $P$ |
| --- | --- | --- | --- | --- | --- | --- |
| MSRA | 8 | 9911830 | 10286401 | **$7.51 \times 10^{-7}$** | 0.002 | **$5.86 \times 10^{-20}$** |
| ZFPM2 | 8 | 106331147 | 106816767 | **$9.12 \times 10^{-7}$** | 0.014 | 0.034 |
| ARNTL | 11 | 13299325 | 13408813 | **$1.07 \times 10^{-6}$** | 0.030 | 0.904 |
| MTMR9 | 8 | 11142000 | 11185655 | **$2.41 \times 10^{-6}$** | $4.69 \times 10^{-4}$ | **$2.50 \times 10^{-12}$** |
| CADM2 | 3 | 85008133 | 86123579 | 0.817 | **$2.75 \times 10^{-7}$** | 0.010 |

Four SNPs, all on chromosome 12, were genome-wide significant for the worry/vulnerability phenotype. These SNPs were located in one locus, spanning 219kb. This region contains the gene *PPFIA2*, which is known to be part of the postsynaptic density in humans^14,15^. Subcomponents of the postsynaptic density have been associated with individual differences in intelligecne^16^ schizophrenia^17^ and mutations in the postsynaptic density have been linked to over 100 brain diseases^14^. The protein encoded by *PPFIA2* has been shown to bind with the calcium-calmodulin-dependent serine protein kinase, a part of the MAGUK family, which appears to be involved in the regulation of higher-order brain function in the mammalian line, again indicating a role for this gene in intelligence, and mental disorders. The association with variation in *PPFIA2* appears to be specific to the factor of worry/vulnerability, as it has not been found in large GWASs conducted on neuroticism^2,3^, although the four SNPs that were genome-wide significant for worry/vulnerability were nominally significant for neuroticism (Table 2.).

Two SNPs were genome-wide significant for the anxiety/tension factor. One was on chromosome 18 (Affymetrix ID 18:27180057_GA_G, Beta = 0.48, *P* = 2.28 × 10^−8^), and the other on chromosome 19 (rs550621052, Beta = 0.43, *P* = 1.32 × 10^−8^). However, these two variants were of low frequency (each MAF = 0.001) and so most likely represent false positives arising from a lack of power to detect association at this MAF.

With the exception of rs2678897 on chromosome 9, the most significant SNPs in each of the independent clumps for the general factor of neuroticism were found to be eQTLs that are involved in the expression of cortical tissue (**Supplementary Table 3**). For the worry/vulnerability factor of neuroticism a significant association was found between rs4360772 and expression changes in the intralobular white matter (*P* = 1.90 × 10^−3^) indicating that rs4360772 is an eQTL that can affect expression differences in the brain in addition to its association with worry/vulnerability.

In order to further investigate the degree to which these three neuroticism phenotypes were genetically distinct, gene-based analysis was conducted using MAGMA.^18^ Thirty-two genes were statistically significant for the general factor of neuroticism; one was significant for the anxiety/tension factor; and four were found for the worry/vulnerability phenotype (**Supplementary Tables 4-6, Table 3**). For the worry/vulnerability phenotype, two of the genome-wide significant genes, *ZFPM2*, and *ARNTL*, did not attain statistical significance in the other two neuroticism variables. Variants in the region of *ZFPM2* have been associated with traits including vascular endothelial growth factor^19^, and variants in the region of *ARNTL* are associated with chronotype^20^, and with body mass index (BMI)^21^. The most significant gene for the anxious/worry factor was *CADM2*, which has previously been associated with processing speed in humans where an intronic variant, rs17518584, attained genome-wide significance in a meta-analysis of ~35,000 individuals. Variants within *CADM2* have also been associated with education, ^22,23^ and BMI^21^.

Partitioning the genome into functional categories indicated more differences between these three phenotypes. Whereas enrichment was found for both the general neuroticism factor and the anxiety/tension factor in regions of the genome that have undergone purifying selection, no such enrichment was found for the worry/vulnerability factor. The largest difference in the pattern of enrichment found was identified when examining which tissues showed enrichment. For each of the three neuroticism phenotypes, significant enrichment was found for the tissues of the central nervous system (general factor fold enrichment = 2.76, *P* = 1.35 × 10^−4^, anxiety/tension fold enrichment = 3.13, *P* = 1.90 × 10^−4^, worry/vulnerability fold enrichment = 3.57, *P* = 2.79 × 10^−4^); however, for the anxiety/tension factor significant enrichment was also found for the adrenal/pancreas (fold enrichment = 4.57, *P* = 6.52 × 10^−4^), cardiovascular (fold enrichment = 3.76, *P* = 0.004), and skeletal/muscle tissues (fold enrichment = 3.14, *P* = 0.007) (**Supplementary Table 7, Figure 2**).

Next, we examined the overlap between the genetic architecture of the three neuroticism traits with health, anthropometric, SES, longevity, reproductive, and wellbeing variables. Alzheimer’s disease was run with and without the *APOE* region in order to prevent the large associations in this region biasing the regression model. Full details of the GWAS that provided summary statistics for each of the 31 phenotypes are provided in Supplementary Table 8.

Following false discovery rate (FDR) correction for multiple comparisons, 16 of the 32 genetic correlations were statistically significant for the general factor of neuroticism, 16 of the 32 were significant for the anxiety/tension factor, and 14 of the 32 were statistically significant for the worry/vulnerability factor. (**Figure 3**. and **Supplementary Table 9**.). Whereas the number of genetic correlations was similar between these traits, in many instances the direction of effect is reversed from those studies which have found that high levels of neuroticism are associated with poor health, lower cognitive ability, and a lower level of socio-economic status.

**Figure 2.**
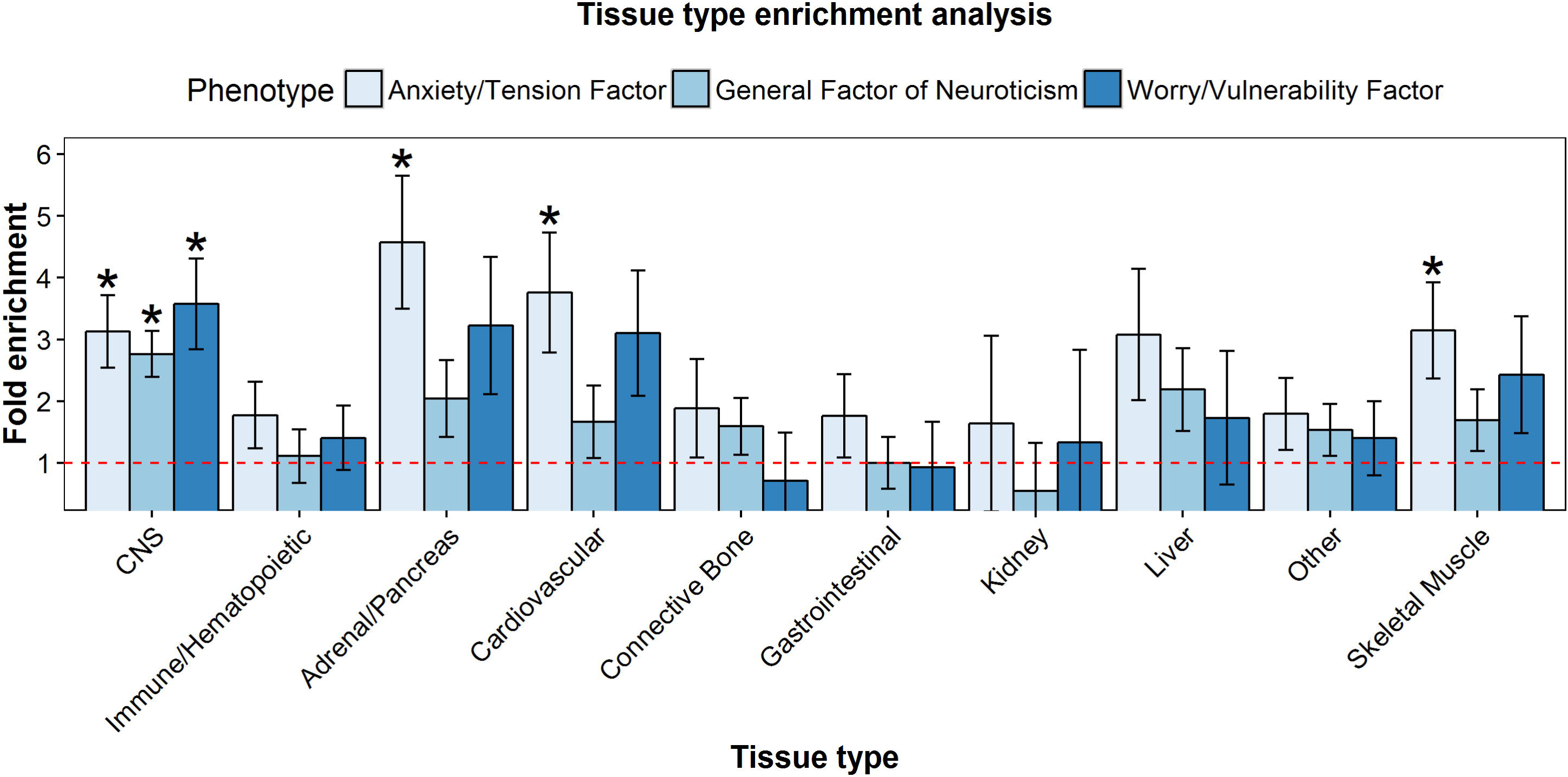
Tissue enrichment for the general factor of neuroticism, the anxiety/tension factor, and the worry vulnerability factor. The red line indicates no fold enrichment and asterisk indicate statistical significance following FDR correction.

**Figure 3.**
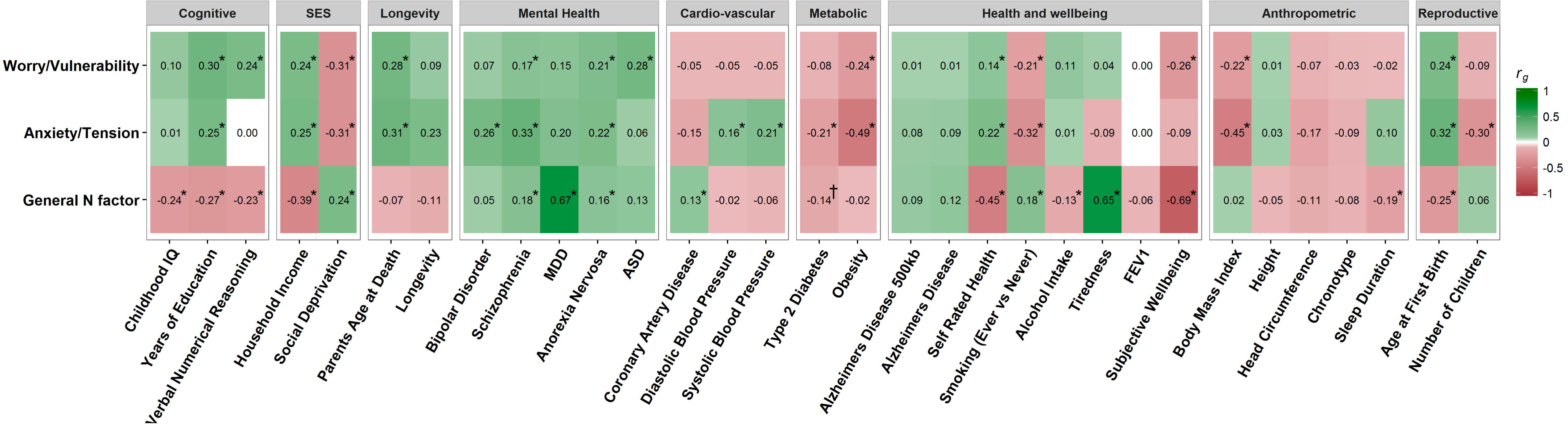
Genetic correlations between the general factor of neuroticism, the anxiety/tension factor, the worry/vulnerability factor with 31 cognitive/socioeconomic/health traits. Colour indicates the direction of the correlation and shade indicates the magnitude of the correlation. Asterisk indicates statistical significance after controlling for 32 tests using FDR, and a dagger indicates nominal significance that did not withstand FDR correction.

For the cognitive variables, the general factor of neuroticism showed significant negative genetic correlations with childhood IQ (*r*_*g*_ = −0.24, *P* = 2.61 × 10^−3^), years of education (*r*_*g*_ = −0.27, *P* = 5.07 × 10^−15^), and with verbal numerical reasoning (*r*_*g*_ = −0.23, *P* = 7.95 × 10^−4^). However, the worry/vulnerability factor showed significant positive genetic correlations with both years of education (*r*_*g*_ = 0.30, *P* = 4.18 × 10^−16^), and with verbal numerical reasoning (*r*_*g*_ = 0.24, *P* = 2.45 × 10^−4^). The anxiety/tension factor also displayed a positive genetic correlation with years of education (*r*_*g*_ = 0.25, *P* = 5.17 × 10^−10^). The genetic correlations with the socioeconomic status variables also showed the same evidence that neuroticism is a composite phenotype, composed of factors with different predictive capabilities. The genetic variants associated with an increase in the general factor of neuroticism were also associated with a genetic risk for a lower household income (*r*_*g*_ = −0.39, *P* = 2.67 × 10^−16^), and living in an area with a higher level of social deprivation (*r*_*g*_ = 0.24, *P* = 6.95 × 10^−5^). However, both the anxiety/tension, and the worry/vulnerability factors showed significant genetic correlations in the opposite directions to the general factor of neuroticism for both household income (anxiety/tension *r*_*g*_ = 0.25, *P* = 7.64 × 10^−4^, worry/vulnerability *r*_*g*_ = 0.24, *P* = 3.57 × 10^−4^), and living in an area with a higher level of social deprivation (anxiety/tension *r*_*g*_ = −0.31, *P* = 3.87 × 10^−5^, worry/vulnerability *r*_*g*_ = −0.31, *P* = 5.02 × 10^−5^). This indicates that, whereas neuroticism may be associated with a lower socioeconomic status, the two factors measured here appear to be associated with advantages that, in turn, are associated with the acquisition of wealth and improved living conditions.

This difference between the genetic correlations derived using the general factor and the worry/vulnerability, and the anxiety/tension factors was also found for age at first birth; the anxiety/tension and worry/vulnerability factors were genetically correlated with delaying childbirth (anxiety/tension *r*_*g*_ = 0.32, *P* = 7.87 × 10^−9^, worry/vulnerability *r*_*g*_ = 0.24, *P* = 2.23 × 10^−6^). On the other hand, the negative genetic correlation (*r*_*g*_ = −0.25, *P* = 8.12 × 10^−7^) found between the general factor of neuroticism and the age of first birth indicates that the genetic variants that are associated with increases in neuroticism are also associated with a lower age at which an individual has a child. Self-rated health and smoking also showed this pattern of results whereby significant genetic correlations were identified for the three traits but with the opposite direction of effect between the general factor and the two factors. For the general factor a positive genetic correlation was found with smoking (*r*_*g*_ = 0.18, *P* = 3.91 × 10^−4^), and a negative genetic correlation was found with self-rated health (*r*_*g*_ = −0.45, *P* = 2.88 × 10^−8^). For anxiety/tension, and worry/vulnerability negative genetic correlations were found with smoking, (anxiety/tension, *r*_*g*_ = −0.32, *P* = 1.02 × 10^−5^; worry/vulnerability, *r*_*g*_ = −0.21, *P* = 2.12 × 10^−3^), and positive genetic correlations were found with self-rated health (anxiety/tension, *r*_*g*_ = 0.22, *P* = 1.48 × 10^−3^; worry/vulnerability, *r*_*g*_ = 0.14, *P* = 1.86 × 10^−2^). This pattern of genetic correlations indicates a protective effect of these alleles against smoking in addition to self-perceived better health.

The two neuroticism factors of anxiety/tension and worry/vulnerability also displayed unique genetic correlations not found using the general factor of neuroticism, again indicating that these factors have elements of their genetic architecture that do not overlap with the general factor of neuroticism. Most strikingly, whereas no genetic correlations were found between the general factor of neuroticism and parent’s age at death (*r*_*g*_ = −0.07, *P* = 0.56), a proxy measure for the longevity of the current participants of UK Biobank, significant positive genetic correlations were found with each of the two factors (anxiety/tension *r*_*g*_ = 0.31, *P* = 8.05 × 10^−3^, worry/vulnerability *r*_*g*_ = −0.28, *P* = 1.10 × 10^−2^), indicating that the genetic contribution of these two traits also acts to increase longevity. Genetic correlations were also found for each of the factors—but, importantly, not with the general neuroticism factor—with obesity (anxiety/tension *r*_*g*_ = 0.49, *P* = 8.01 × 10^−9^, worry/vulnerability *r*_*g*_=-0.24, *P* = 6.39 × 10^−7^), and BMI (anxiety/tension *r*_*g*_ = −0.45, *P* = 4.06×10^−27^, worry/vulnerability *r*_*g*_=-0.49, *P* = 8.01 × 10^−22^). These results indicate a protective association against obesity, and a high BMI generally, for variants associated with increasing levels of the anxiety/tension, and worry/vulnerability factors. However, in the case of the anxiety/tension factor a positive genetic correlation was found with both systolic and diastolic blood pressure (*r*_*g*_ = 0.21, *P* = 2.19×10^−3^, *r*_*g*_ = −0.16, *P* = 2.48 × 10^−2^), indicating that the genetic burden for this factor is associated with unique health problems.

Whereas the pattern of significant genetic correlations differs between general factor and the two factors of neuroticism, they each showed significant genetic correlations with mental health, with none of the three phenotypes affording apparent genetic protection against mental illness. Differences between the general factor of neuroticism, and the two factors were still apparent as, whereas each of the three phenotypes showed a positive genetic correlation with schizophrenia (general factor *r*_*g*_ = 0.18, *P* = 2.89 × 10^−4^, anxiety/tension *r*_*g*_ = 0.33, *P* = 7.88 × 10^−12^, worry/vulnerability *r*_*g*_ = 0.17, *P* = 1.09 × 10^−4^), only the general factor showed a significant genetic correlation with major depressive disorder (*r*_*g*_ = 0.67, *P* = 1.64 × 10^−14^). Each of the factors also showed their own unique overlap with mental health, as only the anxiety/tension factor showed a significant genetic correlation with bipolar disorder (*r*_*g*_ = 0.26, *P* = 1.03 × 10^−3^), and only the worry/vulnerability factor showed a significant genetic correlation with autistic spectrum disorder (*r*_*g*_ = 0.28, *P* = 7.97 × 10^−4^). In line with the genetic correlations with mental health, the genetic correlations between the general factor of neuroticism, and each of the factors, with subjective well-being was negative, although it was not significant for the anxiety/tension factors (general factor *r*_*g*_ = −0.69, *P* = 2.49 × 10^−49^, anxiety/tension *r*_*g*_ = −0.09, *P* = 0.22, worry/vulnerability *r*_*g*_ = −0.26, *P* = 3.44 × 10^−5^).

Neuroticism is widely recognised as one of the most prominent personality traits, with high importance in causing personal, societal and financial burdens of human misery^9^. However, this new work shows, at the genetic level, that neuroticism is molecular and not atomic; it has parts that are risks for and parts that are protectors of physical health, and longevity. The strongest evidence for this is shown in how the genetic architecture of two factors measured here—anxiety/tension, and worry/vulnerability—show different overlaps with health, cognitive, and socioeconomic status when compared with general neuroticism. A higher polygenic burden for the general factor of neuroticism is an indication for a genetic risk for lower cognitive ability, lower SES, an increase in coronary artery disease, and lower self-rated health. However, the polygenic contribution to two orthogonal factors of neuroticism—worry/vulnerability and anxiety/tension— is associated with higher levels of cognitive ability, more affluent socioeconomic status, a perceived better level of overall health, and longer life.

## Acknowledgements

This work was undertaken in The University of Edinburgh Centre for Cognitive Ageing and Cognitive Epidemiology (CCACE), supported by the cross-council Lifelong Health and Wellbeing initiative (MR/K026992/1). Funding from the Biotechnology and Biological Sciences Research Council (BBSRC), the Medical Research Council (MRC), and the University of Edinburgh and gratefully acknowledged. CCACE funding supports IJD. We thank Stuart J Ritchie for assistance with figures. AMM and IJD acknowledge the support of the Wellcome Trust (Wellcome Trust Strategic Award “STratifying Resilience and Depression Longitudinally” (STRADL) Reference 104036/Z/14/Z). We are grateful to have received summary statistics from the International Consortium for Blood Pressure GWAS.

This research has been conducted using the UK Biobank Resource under UK Biobank application 10279

WDH is supported by a grant from Age UK (Disconnected Mind Project).

## Online methods

### Population and study design

Participants were members of the UK Biobank study described in detail elsewhere (http://www.ukbiobank.ac.uk)^24^. In brief UK Biobank consists of 502,655 participants who were recruited between the years of 2006 and 2010 from the United (target age range 40-69 years). Each participant provided detailed information pertaining to their background, lifestyle, as well as undergoing cognitive and physical testing. Additionally, blood, urine, and saliva sample were provided and stored. In the current study we make use of the first release of the genetic data from UK Biobank consisting of 112,151 individuals (52.53% female) aged between 40 and 73 years (mean age = 56.9 years, SD = 7.9) following quality control described below.

### Phenotype measurement

Neuroticism was assessed using 12-questions from the short form of the Revised Eysenck Personality Questionnaire^13^, e.g., “Does your mood often go up and down?”. UK Biobank participants were administered these items using touchscreens and were instructed to “Work quickly and do not think about the exact meaning of the question.” Participants were asked to choose one of four responses for each question: “Yes” (coded 1), “No” (coded 0), “Do not know” (coded −1) and “Prefer not to answer” (coded −3). Of the 502,655 UK Biobank participants, 100,960 did not answer “Yes” or “No” to all 12 questions, and were excluded from further analyses.

We used the data on the remaining 401,695 participants to generate three scores. These three represented the general factor of neuroticism and two orthogonal residual latent traits (also termed specific factors^25^) derived from a bifactor model of neuroticism estimated using exploratory structural equation modelling with an oblique bi-factor Geomin rotation^26,27^. This analysis was carried out in Mplus version 7.4^28^. This approach enabled us to obtain factor scores that were not correlated with the general factor of neuroticism. As such, relationships between lower-order factors and outcomes were not confounded by the associations of these factors with the higher-order general factor^29^.

The bifactor model is presented in Supplementary Table 1. The first factor was primarily defined by three items that indicated how tense or nervous a participant reported themselves to be and was most clearly defined by the question “Would you call yourself a nervous person?” We named this ‘anxiety/tension’. The second factor was primarily defined by three items that indicated how worried or vulnerable a participant reported themselves to be and was most clearly defined by the question “Do you worry too long after an embarrassing experience?” We named this ‘worry/vulnerability’. The correlations between the general factor and the two factors in the model were defined as zero; the correlation between the factors in the model was r = 0.311 (SE = 0.006, 95% CI = [0.301, 0.321], p < 0.0001).

### Genotyping and quality control

A total of 152,729 blood samples were submitted to UK Biobank and were genotyped on either the UKBileve array (N = 49,979) or the UK Biobank axiom array (N = 102,750). Genotyping was performed by Affymetrix on batches of ~4,700 samples. Details of the sample processing specific to UK Biobank can be found at http://biobank.ctsu.ox.ac.uk/crystal/refer.cgi?id=155583, and details pertaining to the Axiom array can be found at http://media.affymetrix.com/support/downloads/manuals/axiom_2_assay_auto_workflow_user_guide.pdf. A stringent quality control procedure was applied by the Wellcome Trust Centre for Human Genetics (WTCHG) to the UK Biobank data prior to its release. Full details of the quality control procedure used can be found at http://biobank.ctsu.ox.ac.uk/crystal/refer.cgi?id=155580.

Additional quality control was also conducted for the current study where individuals were removed based on non-British ancestry (within those who self-identified as being British, principal component analysis was used to remove outliers, N = 32,484), relatedness (N = 7,948), high missingness (N=0), QC failure in UK Bileve (N = 187), as well as gender mismatch (N=0). The final sample size for those with genetic data available was 112,151.

Genome-wide association analysis (GWAS) in the UK Biobank sample

The UK Biobank genetic data was imputed to a reference set that combined the 1000 genomes Phase 3 and the UK10K haplotype reference panels. The full details of this procedure can be found at http://biobank.ctsu.ox.ac.uk/crystal/refer.cgi?id=157020. Following association analysis the data were filtered to exclude SNPs that had a MAF of 0.001 (0.1%) and SNPs with an imputation quality score of <0.1. Following these steps ~17.3 million SNPs remained.

Curation of summary data from GWAS on cognitive, mental health, and anthropometric traits

Genetic correlations were derived between the three neuroticism phenotypes, as well as with 31 traits which show phenotypic correlations with neuroticism and other measures of personality. Full details of each of these GWAS along with links (where possible) to the data used can be found in Supplementary Table 8.

## Statistical analysis

### Genome-wide SNP-based heritability

The total phenotype variance explained by additive genetic effects for each of the three neuroticism phenotypes was quantified using GREML-SC, carried out in GCTA, performed on the genotyped data^30,31^. All genotyped autosomal variants were included in the GREML analysis for the general factor of neuroticism, the anxiety/tension, and the worry/vulnerability factors.

### SNP—based association testing

Single SNP association was carried out using SNPTEST V2.5 (available at https://mathgen.stats.ox.ac.uk/genetics_software/snptest/snptest.html#introduction). Prior to association each of the three phenotypes was adjusted to control for the effects of age, sex assessment centre, genotyping batch, genotyping array, as well as population stratification (14 components). A total of 91,469 participants had both information pertaining to the neuroticism phenotype and genetic data available for association analysis. An additive model was specified using the ‘frequentist1’ option and genotype dosage scores were used in order to account for uncertainty in imputation from the genotyped data. The presence of residual stratification in the GWAS summary statistics was quantified using the intercept of a univariate linkage disequilibrium score (LDSC) regression analysis conducted on each of the three neuroticism phenotypes.

### Clumping

In order to examine the independent regions of the genome tagged by the SNPs in the association analysis linkage disequilibrium (LD) clumping was used. The European panel of the 1000 genomes (phase1, release 3) was used to model LD between the SNPs. Index SNPs were defined as those that reached genome wide significance (5 × 10^−8^), SNPs were included in the clumps if they had a p-value of <1 × 10^−5^, in LD of r^2^ > 0.1, and within 500kb of the index SNP.

### Cortical eQTL analysis

The web resource Braineac was used to examine evidence that the most significant SNPs in each independent region had significant eQTL associations. Braineac uses data from UK Brain Expression Consortium (UKBEC) database containing post-mortem brains from 134 individuals who are free from any known neurological and neurodegenerative disorder.^32^ A total of 10 cortical regions were examined

### Gene-based association analysis

Gene-based analysis was conducted using MAGMA. SNPs from the summary statistics from each of the three neuroticism phenotypes were matched to genes according to the NCBI 37.1 build with gene boundaries being defined as the stop and start site. To model linkage disequilibrium the reference panel from the 1000 genomes (phase 1, release 3) was used. This led to 18 062 autosomal genes being available for analysis. To control for multiple testing a Bonferroni correction was used resulting in an alpha level of 2.77 × 10^−6^ for the three phenotypes.

### Partitioned heritability

Partitioned heritability was conducted using LDSC regression. Firstly, a baseline model was constructed using a total of 25 (arranged into 52 overlapping groups including the original annotations with an additional boundary) functional annotations was used. Secondly, a tissue specific analysis was carried out by including one of the ten tissue types to the baseline model. The goal of this analysis was to determine if any of the tissue types examined were enriched for their contribution to the three neuroticism phenotypes. Enrichment of the partitioned regions was derived separately for each of the three neuroticism phenotype.

